# Comprehensive transcriptomic analysis of cell lines as models of primary tumor samples across 22 tumor types

**DOI:** 10.1101/422592

**Authors:** K. Yu, B. Chen, D. Aran, J. Charalel, A. Butte, T. Goldstein, M. Sirota

## Abstract

Cancer cell lines are commonly used as models for cancer biology. While they are limited in their ability to capture complex interactions between tumors and their surrounding environment, they are a cornerstone of cancer research and many important findings have been discovered utilizing cell line models. Not all cell lines are appropriate models of primary tumors, however, which may contribute to the difficulty in translating *in vitro* findings to patients. Previous studies have leveraged public datasets to evaluate cell lines as models of primary tumors, but they have been limited in scope to specific tumor types and typically ignore the presence of tumor infiltrating cells in the primary tumor samples. We present here a comprehensive pan-cancer analysis utilizing approximately 9,000 transcriptomic profiles from The Cancer Genome Atlas and the Cancer Cell Line Encyclopedia to evaluate cell lines as models of primary tumors across 22 different tumor types. After adjusting for tumor purity in the primary tumor samples, we performed correlation analysis and differential gene expression analysis between the primary tumor samples and cell lines. We found that cell-cycle pathways are consistently upregulated in cell lines, while no pathways are consistently upregulated across the primary tumor samples. In a case study, we compared colorectal cancer cell lines with primary tumor samples across the colorectal subtypes and identified three colorectal cell lines that were derived from fibroblasts rather than tumor epithelial cells. Lastly, we propose a new set of cell lines panel, the TCGA-110, which contains the most representative cell lines from 22 different tumor types as a more comprehensive and informative alternative to the NCI-60 panel. Our analysis of the other tumor types are available in our web app (http://comphealth.ucsf.edu/TCGA110) as a resource to the cancer research community, and we hope it will allow researchers to select more appropriate cell line models and increase the translatability of *in vitro* findings.

## Introduction

Cancer cell lines are an integral part of cancer research and are routinely used to study cancer biology and to screen anti-tumor compounds. While they are relatively inexpensive and easy to grow under laboratory conditions, cell lines have known limitations as preclinical models of cancer and many promising candidate drug compounds have failed to show utility among patient populations^1,2^. Prior studies in ovarian cancer^3^, liver cancer^4^, and breast cancer^5,6^ have shown that cell lines differ in their ability to represent the primary tumors they were derived from, suggesting that using more appropriate cell lines for cancer studies may increase the translatability of preclinical findings. While these previous studies are valuable resources for researchers studying these select tumor types, there is a need for a comprehensive pan-cancer analysis of cell lines and primary tumors.

The generation of large public molecular datasets has allowed researchers to investigate cancer biology at a scale that was unheard of a decade ago. In particular, The Cancer Genome Atlas (TCGA)^7^ research group has collected and characterized the molecular profiles of tumors from over 11,000 patients across 33 different tumor types. They provide clinical, transcriptomic, methylation, copy number, mutation, and proteomic data to facilitate the in-depth interrogation of cancer biology at multiple molecular and clinical levels. Additionally, the Broad Institute’s Cancer Cell Line Encyclopedia^8^ is another large-scale research effort which characterized over 1,000 human-derived cancer cell lines across 36 tumor types and provides transcriptomic, copy number, and mutation data.

Previous studies have integrated data from both of these datasets to evaluate cell lines as models of specific tumor types. For example, Domcke et al. focused primarily on copy number alterations and mutation data to evaluate cell lines as models of high grade serous ovarian carcinomas (HGSOC)^3^. They created a cell line suitability score using features of HGSOC and discovered that the most commonly used cell lines do not seem to resemble HGSOC tumors and the cell lines most representative of HGSOC have very few publications. Similarly, Chen et al. compared hepatocellular carcinoma primary tumor samples to cell lines using transcriptomic data and found that nearly half of the hepatocellular carcinoma cell lines in CCLE do not resemble their primary tumors^4^. In breast cancer, Jiang et al. compared gene expression, copy number alterations, mutations, and protein expression between cell lines and primary tumor samples^5^. They created another cell line suitability score by summing the correlations across all four molecular profiles, although it is notable that only gene expression and copy number alterations had a substantial effect on their score as mutations and protein expression had extremely low correlations across all cell lines (R < 0.1). In another breast cancer study, Vincent et al. compared transcriptomic data between cell lines and primary tumor samples and identified basal and luminal cell lines that were most similar to their respective breast cancer subtypes^6^. While these studies provide insight into specific tumor types, here we hope to provide researchers with a pan-cancer resource that is, to the best of our knowledge, the most comprehensive to date. Additionally, unlike previous studies, we correct for batch effects between different datasets and we adjust for tumor purity which can be a significant confounder in primary tumor transcriptomic data^22^.

The National Cancer Institute’s NCI-60 cell lines are perhaps the most well studied human cancer cell lines and have been in use for nearly three decades by both academic and industrial institutions for drug discovery and cancer biology research^9^. The NCI-60 panel contains 60 human tumor cell lines representing nine human tumor types: leukemia, colon, lung, central nervous system, renal, melanoma, ovarian, breast and prostate. Over 100,000 anti-tumor compounds have been screened using this cell line panel, generating the largest cancer pharmacology database worldwide. While this cell line panel has provided valuable insight into mechanisms of drug response and cancer biology, new large public molecular datasets allow us to compare the NCI-60 cell lines to primary tumor samples and propose more representative cell lines for an improved cancer cell line panel.

In this study, we compared transcriptomic profiles from cell lines and primary tumor samples across the 22 tumor types covered by both TCGA and CCLE. We observed the confounding effect of primary tumor sample purity in our analysis and we adjusted for purity in our subsequent correlation analysis and differential expression analysis of cell lines and primary tumor samples. We found that cell-cycle related pathways are consistently upregulated in cell lines while no pathways are consistently upregulated across the primary tumor samples. We then present our analysis of colorectal adenocarcinoma (COADREAD) cell lines and primary tumor samples and show that we are able to identify cell lines that originated from a different cell type lineage compared to the primary tumor samples. Although only our COADREAD analysis is presented in the main text, we also analyzed the other 21 tumor types and present our results as a web application and a resource to the cancer research community (http://comphealth.ucsf.edu/TCGA110). Lastly, we selected the cell lines that were the most correlated to their primary tumor samples across the 22 tumor types and propose a new cell line panel, the TCGA-110, as a more appropriate and comprehensive panel for pan-cancer studies.

## Results

### Pan-cancer comparison of expression data between cell lines and primary tumor samples

We compared RNA-seq profiles from 9,111 primary tumors from TCGA with 666 cell lines from CCLE across 22 overlapping tumor types (Supplementary Table 1). We normalized counts using the upper-quartile method and corrected for batch effects from the different data sources using the RUVseq package^10^ (Supplementary Figure 1A). We then adjusted for tumor purity in the primary tumor samples and calculated correlation coefficients between primary tumor samples and cell lines using the 5000 most variable genes, as these genes are the most likely to be biologically informative (see Methods).

We found that the median correlation coefficients between cell lines and their matched tumor samples ranged from 0.75 to 0.50 (Figure 1B). Diffuse Large B-cell Lymphoma (DLBC) and Acute Myeloid Leukemia (LAML) have the highest median correlation coefficients of 0.75 and 0.74, suggesting that the blood cancer cell lines are relatively good models of cancer for their primary tumor samples. On the other end of the spectrum, thyroid carcinoma (THCA) and cholangiocarcinoma (CHOL) have the lowest median correlation coefficients of 0.51 and 0.50. Previous studies have shown that the expression profiles of 6 common THCA cell lines (3 of which are included in our study) are significantly more correlated with dedifferentiated anaplastic thyroid carcinoma than the more common papillary thyroid carcinomas^11^, which was the tumor type studied by the TCGA. The low correlations in this study are likely driven by the lack of dedifferentiated anaplastic thyroid carcinoma samples in our primary tumor dataset. Additionally, many of the most common cancers, such as breast invasive carcinoma (BRCA), colorectal adenocarcinomas (COADREAD), and skin cutaneous melanoma (SKCM) have a wide range of correlation coefficients (respectively, R = 0.18-0.78, 0.11-0.86, 0.20-0.85). These ranges likely reflect the amount of heterogeneity within each tumor type and suggest that some primary tumor samples are well matched with cell lines while others may lack representative cell line models.

**Figure 1.**
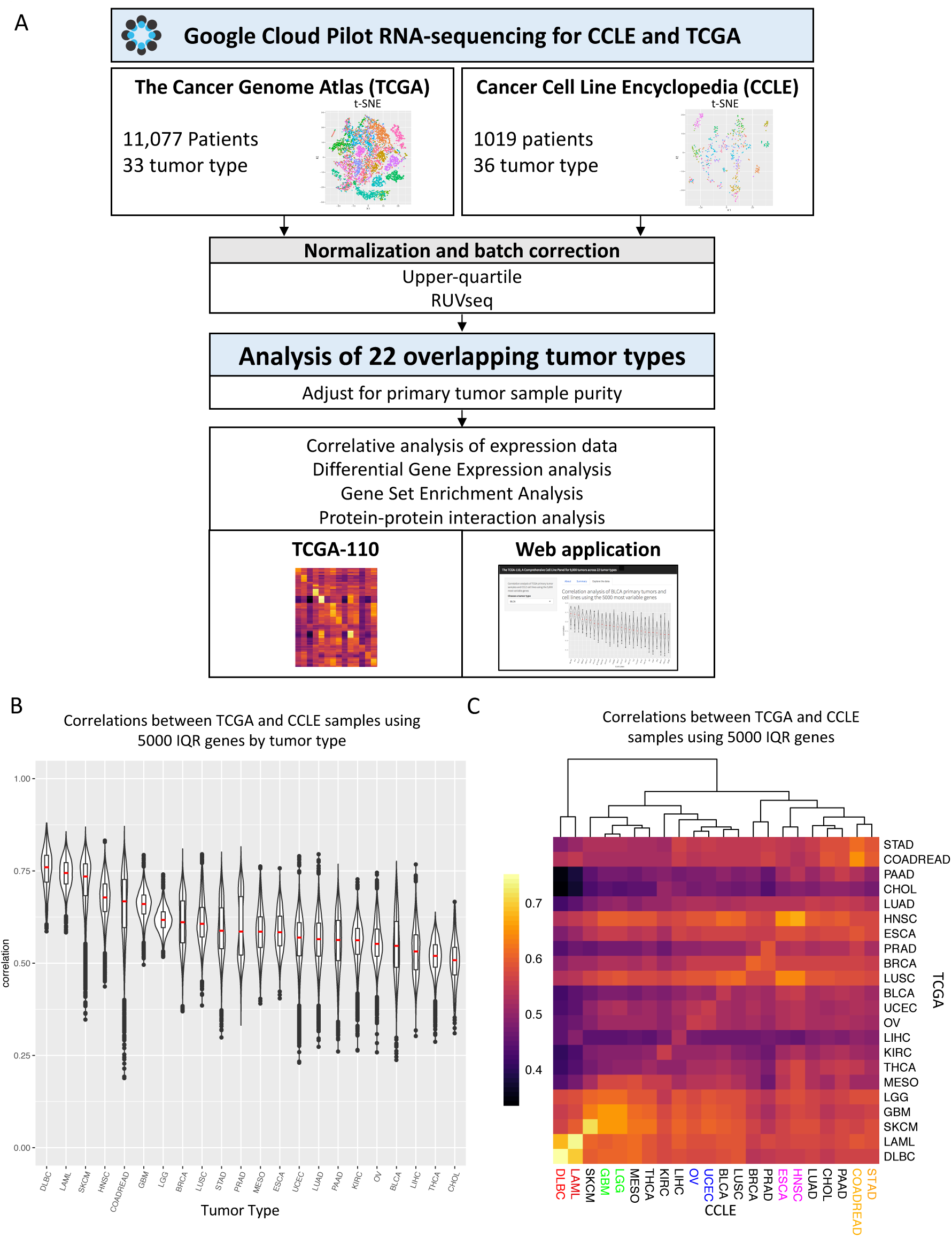
Pan-cancer analysis of cell lines and matching primary tumor samples. A. Study Design. RNA-seq data was downloaded from the Google Cloud Pilot RNA-sequencing for CCLE and TCGA [1] project for 22 cancer types that overlapped between the CCLE and TCGA datasets. The data was normalized, batch corrected, and adjusted for tumor purity for the analysis. B. Correlation analysis of the CCLE and TCGA data. Each sample in the violin plot corresponds to the Spearman correlation between one cell line and one primary tumor sample using the 5,000 most variable genes. C. Heatmap showing the Spearman correlations between cell lines and primary tumor samples across all 22 tumor types summarized by the geometric means. Biologically similar tumor types cluster together (colored text).

Our clustering analysis of cell line and primary tumors correlation coefficients largely captures known biological relationships between the tumor types (Figure 1C). The first split in our clustering analysis depicts the large difference between hematopoietic tumor types and solid tumor types previously shown in other studies^12^. Within the solid tumor cluster, tumor types from similar cell of origin generally clustered together such as ovarian serous cystadenocarcinoma (OV) and Uterine Corpus Endometrial Carcinoma (UCEC), colorectal carcinoma (COADREAD) and stomach adenocarcinoma (STAD), glioblastoma (GBM) and lower grade glioma (LGG), and esophageal carcinoma (ESCA) and head and neck squamous cell carcinoma (HNSC). Interestingly, we observe that sometimes the highest correlation coefficients are not necessarily between cell line and primary tumor samples from the same tumor type. In fact, in 6/22 tumor types, primary tumor samples have higher correlation coefficients with other tumor cell lines than their own. This may indicate poor differentiation in the cell lines or primary tumor sample, lack of appropriate cell line models, or similar expression profiles among certain tumor types (e.g. STAD has the highest correlation with COADREAD).

### Differences between primary tumor sample and cell lines are largely driven by tumor purity

To explore the differences between cell lines and primary tumor samples, we initially performed our correlation and differential gene expression analysis across all 22 tumor types without accounting for tumor purity of the primary tumor samples (Figure 2A and Supplementary Table 2). In our correlation analysis, we compared the cell line correlations with primary tumor samples in the top quartile of tumor purity to the cell line correlations with primary tumor samples in the bottom quartile of tumor purity for each tumor type (Figure 2A). In 19/22 tumor types, the cell lines were significantly more correlated with primary tumor samples in the top quartile of purity, suggesting that the individual correlation coefficients are reflecting, to a certain extent, the degree of tumor infiltration present in the primary tumor samples. Similarly, we found a significant positive relationship (R=0.19, p-value < 2.2e-16) between primary tumor sample purity and the cell line-primary tumor correlation coefficients, suggesting that tumor purity is a significant confounder in our correlation analysis. Furthermore, when we performed Gene Set Enrichment Analysis (GSEA) on the differential expression results using the hallmark gene sets from the MSigDB Collections^13^ and the hallmarks of cancer pathways^14^, we saw that the gene sets involved in immune processes are consistently upregulated in primary tumor samples, suggesting that the largest biological signal from the TCGA samples can likely be attributed to the immune cell infiltrate that are present in the primary tumor samples and absent in the pure cell line populations (Supplementary fig. 2A-2B).

**Figure 2.**
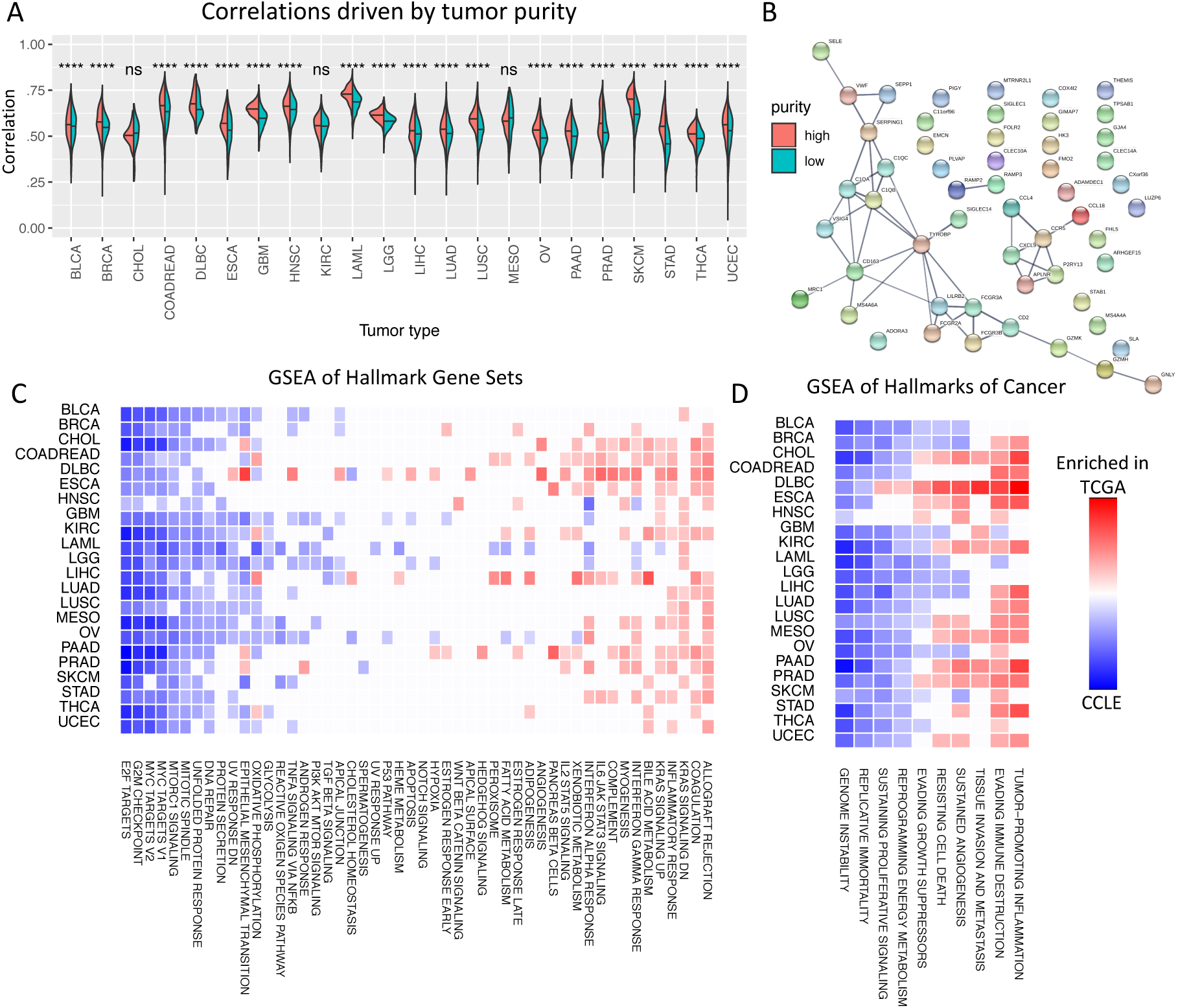
Primary tumor sample/cell line correlations driven by tumor purity. A. Correlations between cell lines and high purity primary tumor samples (red) are significantly higher than correlations between cell lines and low purity primary tumor samples (turquoise) in 19/22 tumor types, motivating our adjustment for tumor purity in subsequent analysis. B. STRING analysis of protein-protein interactions for the 54 genes upregulated in primary tumor samples in at least 20 (90%) of the analyzed tumor types (PPI enrichment p-value < 1.0e-16). Line thickness denotes confidence of the interaction and only high confidence interactions are shown. The PPI network is enriched for immune response pathway genes (5.51e-06). C. Gene Set Enrichment Analysis (GSEA) of differential expression between primary tumor samples and cell lines in hallmark gene sets from MSigDB. NES are shown for pathways with FDR < 5%. Gene sets related to cell cycle progression are enriched in cell lines across all tumor types. D. GSEA of hallmarks of cancer pathways. Genome instability is enriched in cell lines across all tumor types.

After adjusting for primary tumor sample purity in our correlation analysis, we confirmed that there was no longer a significant positive relationship between primary tumor sample purity and cell line-primary tumor correlation coefficients (R=-0.02, p-value < 2.2e-16). Additionally, we found that only 4/22 tumor types retain significantly higher correlations between cell lines and the primary tumor samples in the top quartile of purity compared to cell lines and primary tumor samples in the bottom quartile of purity (Supplementary fig. 2C). We then performed differential expression analysis using tumor purity as a covariate to explore differences in cancer cell biology while minimizing the influence of tumor infiltrating cells. The number of differentially expressed genes ranged from 1,944 in Stomach Adenocarcinoma (STAD) to 5,073 in Low Grade Glioma (LGG) (Supplementary Table 3). We identified 54 genes that were upregulated in primary tumor samples across at least 20 (90%) of the tumor types analyzed and we found a significant number of interactions among these genes (PPI enrichment p-value < 1.0e-16)(Figure 2B). This PPI network was enriched for genes in the immune response pathway (5.51e-06), suggesting that we were not fully able to remove the contribution of the immune infiltrate. However, the GSEA results show a much weaker enrichment of immunological pathways upregulated in the primary tumor samples (Figure 2C-D). Additionally, no pathways were significantly upregulated in primary tumor samples across all the tumor types after adjusting for tumor purity.

No individual genes were significantly upregulated in cell lines across 90% of the tumor types analyzed. However, gene sets involved in cell cycle progression (e.g. E2F targets, G2M checkpoint, Myc targets) and genome instability were significantly enriched in cell lines across all 22 tumor types in our GSEA of MSigDB Hallmark Gene Sets and the Hallmarks of Cancer pathways (Figure 2C-D). These results demonstrate how GSEA can be more informative than analyzing individual upregulated genes alone. Additionally, the enrichment of proliferative gene sets in cell lines across all the tumor types suggests a common response to *in vitro* culturing conditions. Interestingly, unfolded protein response (UPR) was also enriched in cell lines in 17/22 tumor types in our MSigDB Hallmark Gene Set analysis (Figure 2C). This enrichment may reflect a response to the stress caused by the uncontrolled proliferation of cancer cell lines and it suggests that therapies targeting the UPR may show an increased effect in cell lines compared to primary patient tumors.

### Case study: Colorectal cancer cell lines model transcriptomic profiles of some primary tumor subtypes

Colorectal cancer (COADREAD) is the second leading cause of cancer related deaths in the United States and can generally be grouped into four consensus molecular subtypes^15^. These subtypes are CMS1, which are hypermutated, microsatellite unstable, and have strong immune activation, CMS2, which are marked by WNT and MYC signaling activation, CMS3, which are marked by metabolic dysregulation, and CMS4, which are marked by TFG-beta activation, stromal invasion, and angiogenesis. We utilize these subtypes in our study of COADREAD presented here. While only the analysis for COADREAD is shown, analysis of the other tumor types are available in the supplements (Supplementary figures 5-26). For each tumor type, we adjusted for primary tumor purity and compared the expression profiles of the primary tumor samples to the 932 cell line expression profiles in a correlative analysis. We included tumor subtype information when available.

We compared the correlations between COADREAD primary tumor samples and all 932 cell lines grouped by cell line tissue of origin (Figure 3A). The COADREAD primary tumor samples are most correlated with cell lines originating from the large intestine, which contains all the COADREAD cell lines. The correlation coefficients between COADREAD primary tumor samples and cell lines from the large intestine are significantly higher than the correlation coefficients between COADREAD primary tumor samples and cell lines from the stomach, which is the second most correlated tissue of origin (p-value < 1e-16). This confirms that the COADREAD cell lines are more representative of the COADREAD primary tumor samples than cell lines derived from other tissue types.

**Figure 3.**
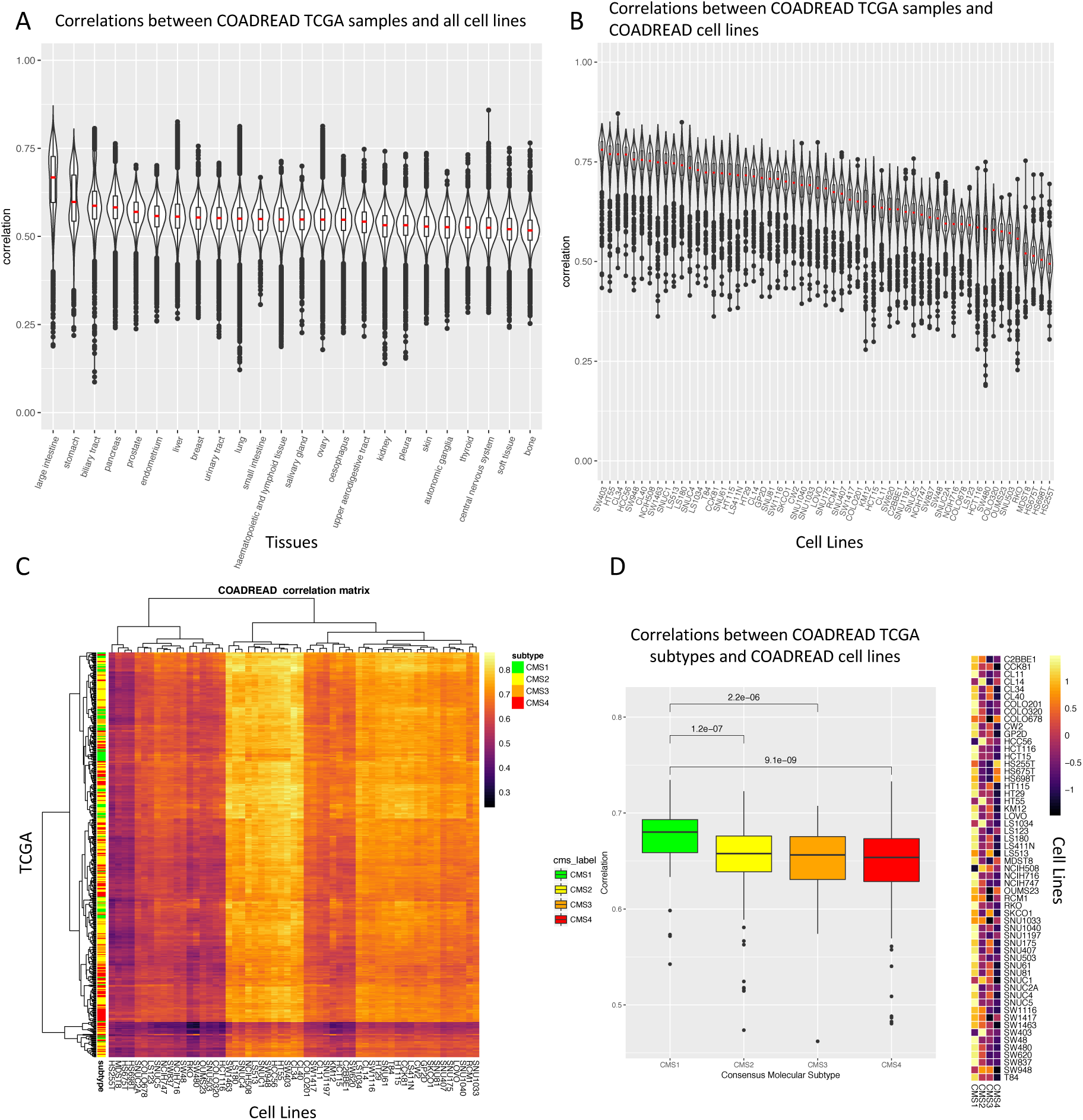
Correlative analysis of colorectal primary tumor samples and cell lines. A. Violin plot of Spearman correlations between primary colorectal tumor samples and all CCLE cell lines using 5,000 most variable genes. The correlations are separated by cell line tissue of origin (x-axis) and the red line is the median correlation coefficient. Primary colorectal tumor samples are significantly more correlated with cell lines from the large intestine (which includes the colon and the rectum) than cell lines from other tissue types (p-value= p-value < 1e-16). B. Correlations between colorectal cell lines and colorectal primary tumor samples, separated by cell lines (x-axis). The median correlation coefficients range from R = 0.76 to R = 0.53. C. Heatmap showing the Spearman correlations between colorectal cell lines (x-axis) and colorectal primary tumor samples (y-axis). The color bar on the y-axis indicate the consensus colorectal molecular subtype [3] of each primary tumor sample. D. Boxplot of correlations between colorectal cell lines and colorectal primary tumor samples separated by the consensus molecular subtypes of the primary tumor samples. Correlations between cell lines and CMS1 primary tumor samples are significantly higher than the other molecular subtypes. Heatmap shows median correlations between cell lines and Consensus Molecular Subtypes, scaled by row.

We next compared individual COADREAD cell lines to the COADREAD primary tumor samples (Figure 3B). The median correlation coefficients of the cell lines ranged from 0.74 to 0.48, suggesting that some cell lines are much more suitable as models of primary tumor samples than others. Within the cell lines, however, the standard deviations of the correlation coefficients are relatively low (0.09 – 0.05). This suggests that between cell line differences are larger than within cell line differences, the latter of which reflects the variability of the primary tumor samples. Interestingly, we found that three cell lines (Hs 675.T, Hs 698.T, Hs 255.T) with significantly lower median correlation coefficients with the primary tumor samples have been annotated as having a fibroblast lineage in the DepMap database. This reveals the strength of the correlative analysis approach, as we were able to identify cell lines derived from tumor microenvironment cells rather than tumor cells.

Next, we incorporated primary tumor subtype information from Guinney et al. which classified the TCGA primary tumor samples into one of four consensus molecular subtypes. Although we did not see strong clustering by primary tumor subtypes in our primary tumor/cell line correlation matrix (Figure 3C), we were able to see significantly higher correlations in CMS1 when we separated the correlation coefficients by molecular subtypes (Figure 3D). Of note, in our initial correlation analysis we did not account for tumor purity and found significantly lower correlations between the cell lines and the CMS4 subtype, which is characterized by stromal invasion (Supplementary Figure 3A). After adjusting for tumor purity, we do not find lower correlations between the cell lines and the CMS4 subtype (Figure 3D), once again illustrating the importance of accounting for the cellular composition of the primary tumor samples

We were able to obtain tumor subtype information for 19/22 tumor types, which we included in our analysis (Supplementary Figures 5-26). Across 18/19 of the tumor types where subtype information was available, we found statistically significant differences between cell line and primary tumor correlations when grouped by primary tumor subtypes. This suggests the importance of considering tumor subtype information when selecting cell lines for cancer studies.

### Molecular data driven approach identifies the improved TCGA-110 cell line panel

The NCI-60 panel of human tumor cell lines has been used in cancer research for over 20 years to screen chemical compounds and natural products. It contains cell lines from the following 10 tumor types: BRCA, COADREAD, GBM, KIRC, LAML, LUAD, LUSC, OV, PRAD, and SKCM. We wanted to determine if the NCI-60 panel could be improved by using cell lines with higher correlations to their primary tumor samples. We analyzed the cell lines that overlapped between the NCI-60 panel and the CCLE database and found that the cell lines in the NCI-60 panel did not have the highest correlations with their primary tumor samples based on gene expression profiles (Figure 4A-C). We created an improved NCI-60 panel by selecting the same number of cell lines per tumor type as the original NCI-60 panel, but choosing the cell lines with the highest correlations per tumor type. The correlations in our improved NCI-60 panel were significantly higher than the original NCI-60 panel, which suggests that the integration of primary tumor data can be used to guide cell line selection for more representative models of cancer.

**Figure 4.**
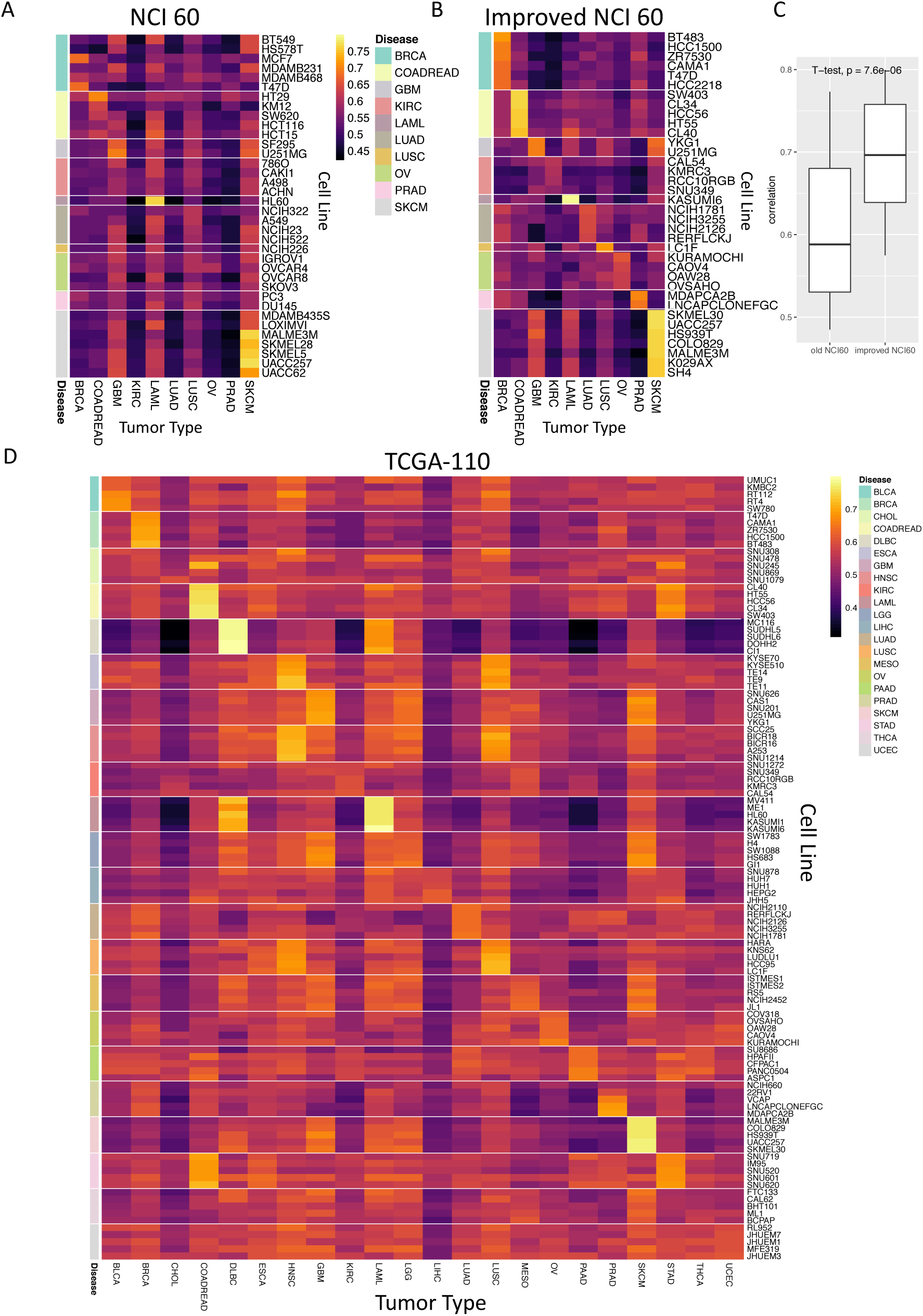
The TCGA-110: an improved cell line panel integrating TCGA and CCLE data. A. Heatmap of correlations between cell lines in the NCI60 panel and primary tumor data (only 36 cell lines which are shared between NCI60 panel and CCLE are shown). Blue boxes indicate cell lines that belong to the tumor type on the x-axis. B. Heatmap of improved NCI60 panel. Improved panel has the same number of cell lines and tumor types as the original NCI60 panel, but the cell lines with the highest correlations with their matched primary tumor samples were selected. Only 3 cell lines overlap between the original NCI60 panel and the improved NCI60 panel. C. Boxplot showing that the improved NCI60 panel has significantly higher correlations with their matched primary tumor samples than the old NCI60 panel. D. Proposed TCGA-110 panel. An improved cell line panel that includes 5 cell lines with the highest correlations to their matched primary tumor samples across 22 tumor types.

We furthermore propose a new expanded panel of cell lines, which we name TCGA-110, to be used as a pan-cancer resource for cancer research and drug screening (Figure 4D, Supplementary Table 4). We selected the 5 cell lines with the highest correlations to their primary tumor samples from each of the 22 tumor types analyzed in this paper to generate our TCGA-110 panel. By using TCGA primary tumor data to guide our cell line selection, we hope that our new panel will be more comprehensive and representative of primary tumor samples than the NCI-60 panel.

## Discussion

While cell lines are commonly used as models of primary tumors in cancer research, cell lines differ from primary tumors in biologically significant ways and not all cell lines may be appropriate models for their annotated tumor type. Previous studies of ovarian cancer, breast cancer, and liver cancer have shown that the molecular profiles of cell lines from the same tumor type can differ widely and some cell lines more closely model their primary tumors than others. In this study, we leveraged publically available transcriptomic data to perform a comprehensive pan-cancer analysis across 22 tumor types and provide a resource for researchers to select appropriate cell lines for their tumor-specific studies.

Our analysis reveals that primary tumor and cell line correlations vary widely across tumor types, with the hematopoietic tumor types having relatively good cell line models and thyroid carcinomas having particularly poor cell line models. Based on previous studies, the thyroid carcinoma cell lines likely model a more dedifferentiated form of resource thyroid carcinomas than the papillary form that was collected for the TCGA study. Clustering tumor types by correlations between primary tumor samples and cell lines generally grouped similar tumor types together. Of note, the primary tumor samples in 6/22 tumor types have higher correlation coefficients with cell lines from other tumor types than cell lines from their own tumor type. These tumor types may contain poorly differentiated samples, which would make it difficult to distinguish them from other tumor types using transcriptomics alone.

We identified primary tumor sample purity as a significant confounder in our correlation and differential expression analysis and show that we are largely able to remove the confounding effect of tumor purity in our analysis. After correcting for primary tumor purity, we found a significantly lower enrichment of immune pathways among the primary tumor samples in our GSEA analysis. While no pathways are consistently upregulated in the primary tumor samples across all the tumor types, we found that cell-cycle related pathways are consistently upregulated in cell lines across all tumor types, perhaps reflecting *in vitro* culturing conditions.

In our case study comparing colorectal cell lines to colorectal primary tumor samples, we found that the colorectal cell lines are more representative of colorectal primary tumor samples than cell lines from other tissues of origin. We also found a wide range of correlation coefficients among the colorectal cell lines, suggesting that the cell lines at the lower end may not be appropriate models of primary colorectal tumors. Indeed, the three colorectal cell lines with the worst correlations with the primary tumor samples seem to be derived from fibroblasts rather than epithelial cells. Lastly, we found that the cell lines had significantly higher correlations with the CMS1 colorectal subtype. While we presented our analysis of colorectal cancer here, we also analyzed the other 21 tumor types and present the results in the supplementary figures (Supplementary Figures 5-26).

Finally, we propose the TCGA-110 cell line panel as a resource for pan-cancer studies. It encompasses 22 different tumor types and contains the cell lines most correlated with their primary tumor samples. We hope that using more representative cell lines will improve our ability to translate cell line findings into patients.

There are several limitations of our study that should be recognized. Although we were not able to match all of the cell lines from CCLE to primary tumor samples in TCGA, we were able to match a majority of the cell lines (71%) to a matching primary tumor type and we provide analysis for less common tumor types whose cell lines have not been well studied. Additionally, despite using samples from different studies collected at different times, we were able to harmonize the data and remove batch effects for our pan-cancer analyses using the RUVseq package. Finally, although our cell line findings lack experimental validation, we were able to identify colorectal cell lines that were derived from fibroblasts rather than tumor epithelial cells, showing the power of our correlative analysis using transcriptomic data.

In future studies, we hope to integrate other types of molecular data such as mutation, copy number alteration, and methylation profiles to provide a multi-omic comparison of cell lines and primary tumor samples.

By leveraging expression profiles from thousands of primary tumor and cell line samples, our study has created a comprehensive pan-cancer resource to aid researchers in selecting the most representative cell line models. We hope that using more appropriate cell line models for cancer studies will allow the research community to better understand cancer biology and translate more *in vitro* findings into clinically relevant therapies.

## Methods

### Data collection and normalization

CCLE cell lines were manually matched to TCGA tumor types using the CCLE Cell Line Annotations file (CCLE_sample_info_file_2012-10-18.txt), which contains histological information for each cell line. While 934 CCLE samples were available in the OSF open-access repository, we were able to match approximately 70% of the samples (n = 679) to their respective TCGA tumor type. We used these matched CCLE cell lines for comparison with TCGA primary tumor samples. These samples encompass the following 22 tumor types: BLCA, BRCA, CHOL, COADREAD, DLBC, ESCA, GBM, LGG, HNSC, KIRC, LAML, LIHC, LUAD, LUSC, MESO, OV, PAAD, PRAD, SKCM, STAD, THCA, UCEC. For the correlative analysis based on cell line tissue of origin, all 934 CCLE samples were used.

TCGA and CCLE RNA-seq samples for the 22 tumor types above were downloaded from the Google Cloud Pilot RNA-Sequencing for CCLE and TCGA project in the OSF open-access repository (https://osf.io/gqrz9/). This repository contains 12,307 RNA-seq samples from both the CCLE and the TCGA databases which have been uniformly processed from raw data.

Transcript alignment and quantification were performed using kallisto (version 0.43.0) and both transcript per million (TPM) values and transcript counts are available in the repository. The transcript counts were downloaded and summarized to the gene-level for this analysis.

To remove batch effects from the two data sources (TCGA and CCLE), we used the RUVg method from the RUVseq^10^ package which uses factor analysis on a set of negative control genes to estimate the factors of unwanted variation. Because a comprehensive set of negative control genes has not been published in cancer, we selected a panel of 1000 empirically derived negative control genes as described in the RUVseq manual to estimate the factors of unwanted variation (Supplementary Figure 1C). RUVseq normalization improved PCA clustering based on tumor type (Supplementary Figure 1A).

We collected tumor purity estimates for the TCGA samples from the following sources: InfiniumPurity^16^, which uses methylation microarray data to estimate tumor purity, ESTIMATE^17^, which uses expression data to estimate purity, ABSOLUTE^18^, which uses relative copy profiles to estimate purity, and IHC^19^, which uses haematoxylin and eosin stain slides produced by the Nationwide Children’s Hospital Biospecimen Core Resource to estimate purity. Since not all samples were analyzed with every purity estimation method, we only used samples for which there were at least two different sources of purity estimates. We summarized the purity estimates for these samples by finding the median purity value and used this for our purity analysis.

### Correlative Analysis

We filtered for lowly expressed genes by removing genes that had fewer than 1 Counts per million (CPM) in at least 90% of the samples. To correct for the heterogeneous cellular composition of the primary tumor samples, we removed genes that have high correlations with tumor purity (R > -0.3, p-value < 0.01) and then adjusted for tumor purity in the primary tumor samples using linear regression. We then selected the top 5000 genes ranked by interquartile range across all TCGA samples for our correlative analysis. We decided to use 5000 genes based on previous studies^4^, although we tried other numbers (1000, 10000) and found our results to be remarkably robust (Supplementary Figure 4A-B).

### Differential Expression and GSEA

We identified differentially expressed genes using the negative binomial GLM approach from the edgeR package^20^. The factors of unwanted variation that were estimated from the RUVseq package were added as a covariate to the design matrix to account for batch effects. We added tumor purity estimates of the primary tumor samples as covariates and we set the tumor purity estimates of all the cell lines as 1. We considered a gene to be differentially expressed if the false discovery rate < 0.01 and the absolute log fold expression change > 2.

For our GSEA analysis, we ranked our genes by their log fold-change values. We then used the GSEAPreanked^21^ software with the “classic” enrichment score, which was recommended for RNA-seq data in the GSEA manual. The enrichment score (ES) reflects the degree to which a gene set is overrepresented at the top or bottom of the ranked list of genes. We downloaded the 50 Hallmark gene sets from the MSigDB Collections^22^ and created our own gmx file for the Hallmarks of cancer pathways using gene sets from the Oncology Models Forum^14^.

## Acknowledgements

Butte, Aran and Goldstein graciously acknowledge the support from the National Cancer Institute Oncology Model Forum project, NIH grant U24 #CA195858. DA is supported by the Gruss Lipper Postdoctoral Fellowship, BC by R21 TR001743 and K01 ES028047, and MS by National Library of Medicine NIH grant K01 LM012381. We would like to thank Boris Oskotsky for technical support and members of the Sirota Lab for useful discussion. We would also like to thank researchers at the Broad Institute and The Cancer Genome Atlas Consortium who released data to the public. The content is solely the responsibility of the authors and does not necessarily represent the official views of the National Institutes of Health.

## References

1 Gillet, J.-P., Varma, S. & Gottesman, M. M. The Clinical Relevance of Cancer Cell Lines. JNCI Journal of the National Cancer Institute 105, 452–458 (2013).

2 Weinstein JN. Nature. 2012 Mar 28; 483(7391):544-5.

3 Domcke, S., Sinha, R., Levine, D. A., Sander, C. & Schultz, N. Evaluating cell lines as tumour models by comparison of genomic profiles. Nature Communications 4, (2013).

4 Chen, B., Sirota, M., Fan-Minogue, H., Hadley, D. & Butte, A. J. Relating hepatocellular carcinoma tumor samples and cell lines using gene expression data in translational research. BMC Medical Genomics 8, (2015).

5 Vincent, K. M., Findlay, S. D. & Postovit, L. M. Assessing breast cancer cell lines as tumour models by comparison of mRNA expression profiles. Breast Cancer Research 17, (2015).

6 Jiang, G. et al. Comprehensive comparison of molecular portraits between cell lines and tumors in breast cancer. BMC Genomics 17, (2016).

7 The Cancer Genome Atlas (TCGA) Research Network. http://cancergenome.nih.gov

8 Broad Institute Cancer Cell Line Encyclopedia. https://portals.broadinstitute.org/ccle

9 Shoemaker, R. H. The NCI60 human tumour cell line anticancer drug screen. Nature Reviews Cancer 6, 813–823 (2006).

10 Risso, D., Ngai, J., Speed, T. P. & Dudoit, S. Normalization of RNA-seq data using factor analysis of control genes or samples. Nature Biotechnology 32, 896–902 (2014).

11 Saiselet, M. et al. Thyroid cancer cell lines: an overview. Frontiers in Endocrinology 3, (2012).

12 Domcke, S., Sinha, R., Levine, D. A., Sander, C. & Schultz, N. Evaluating cell lines as tumour models by comparison of genomic profiles. Nature Communications 4, (2013).

13 Liberzon, A. et al. The Molecular Signatures Database Hallmark Gene Set Collection. Cell Systems 1, 417–425 (2015).

14 Datta, D., Goldstein, T., Gu, Z. & Butte, A. Abstract LB-006: Oncology model fidelity scores. Cancer Research 77, LB-006-LB-006 (2017).

15 Guinney, J. et al. The consensus molecular subtypes of colorectal cancer. Nature Medicine 21, 1350–1356 (2015).

16 Zhang, N. et al. Predicting tumor purity from methylation microarray data. Bioinformatics 31, 3401–3405 (2015).

17 Yoshihara, K. et al. Inferring tumour purity and stromal and immune cell admixture from expression data. Nature Communications 4, (2013).

18 Carter, S. L. et al. Absolute quantification of somatic DNA alterations in human cancer. Nature Biotechnology 30, 413–421 (2012).

19 Aran, D., Sirota, M. & Butte, A. J. Systematic pan-cancer analysis of tumour purity. Nature Communications 6, (2015).

20 Robinson, M. D., McCarthy, D. J. & Smyth, G. K. edgeR: a Bioconductor package for differential expression analysis of digital gene expression data. Bioinformatics 26, 139–140 (2009).

21 Subramanian, A. et al. Gene set enrichment analysis: A knowledge-based approach for interpreting genome-wide expression profiles. Proceedings of the National Academy of Sciences 102, 15545–15550 (2005).

